# Modeling responses of macaque and human retinal ganglion cells to natural images using a convolutional neural network

**DOI:** 10.1101/2024.03.22.586353

**Authors:** Alex R. Gogliettino, Sam Cooler, Ramandeep S. Vilkhu, Nora J. Brackbill, Colleen Rhoades, Eric G. Wu, Alexandra Kling, Alexander Sher, Alan M. Litke, E.J. Chichilnisky

## Abstract

Linear-nonlinear (LN) cascade models provide a simple way to capture retinal ganglion cell (RGC) responses to artificial stimuli such as white noise, but their ability to model responses to natural images is limited. Recently, convolutional neural network (CNN) models have been shown to produce light response predictions that were substantially more accurate than those of a LN model. However, this modeling approach has not yet been applied to responses of macaque or human RGCs to natural images. Here, we train and test a CNN model on responses to natural images of the four numerically dominant RGC types in the macaque and human retina – ON parasol, OFF parasol, ON midget and OFF midget cells. Compared with the LN model, the CNN model provided substantially more accurate response predictions. Linear reconstructions of the visual stimulus were more accurate for CNN compared to LN model-generated responses, relative to reconstructions obtained from the recorded data. These findings demonstrate the effectiveness of a CNN model in capturing light responses of major RGC types in the macaque and human retinas in natural conditions.

## Introduction

A major focus of research on the retina has been to develop models of retinal ganglion cell (RGC) responses to visual stimulation. A commonly used model composed of a spatiotemporal linear filtering stage followed by a rectifying nonlinearity is known as a linear-nonlinear (LN) cascade. This model has been developed and tested primarily on RGC responses to artificial stimuli, such as white noise [1,2]. However, more recently, investigations of the retina have included analysis of responses to more ethologically-relevant stimuli, such as natural images [3–8]. Natural images have very different stimulus statistics from white noise [9], including significant spatiotemporal correlations, which drive RGC firing very differently than white noise does. Indeed, although LN models have been shown to predict RGC responses to white noise with reasonable accuracy [1,2,5], their ability to predict responses to natural images is more limited, indicating that more sophisticated models will be needed to more fully understand how the retina processes natural visual information.

Recent work [6,8] developed a three-layer convolutional neural network (CNN) model to predict responses of salamander RGCs to white noise and natural images. This model predicted RGC responses more accurately than a LN model did and also reproduced several specific nonlinear and potentially behaviorally-relevant retinal computations that have previously been observed experimentally in several species [6]. Furthermore, analysis of the CNN model parameters revealed similarities between the learned convolutional filters and the receptive fields of bipolar cells [6]. More recently, several studies used CNN-based models to capture contrast adaptation in white noise in the macaque and rat retina [10], identify the chromatic properties of mouse RGCs [11] and test how effectively a model trained on a single stimulus type can capture both white noise and natural image responses in the marmoset retina [12]. However, it remains unclear how effectively CNN-based approaches can capture the responses of RGCs in the macaque and human retinas to natural scenes, and the visual information conveyed by those responses.

Here, we apply a CNN light response model to predict responses of the four numerically dominant RGC types in the macaque and human retina – ON parasol, OFF parasol, ON midget and OFF midget – to natural images. We first determined how accurately the CNN model predicts the firing rates of each of these cell types over time, using the LN model as a baseline. We then employed linear image reconstruction to test how accurately model-generated responses capture the visual signal conveyed by the retina. The findings reveal that the CNN model accurately captures responses to natural images of the major RGC types in the macaque and human retina and substantially outperforms the LN model.

## Results

We evaluate the performance of a CNN light response model as follows. We first train and test the model on preparations of three macaque retinas and one human retina, during stimulation with jittered natural images, focusing on ON and OFF parasol and midget cells. We then quantify performance by measuring the response prediction accuracy. Finally, we compare linear reconstructions of the visual stimulus from measured spike responses to reconstructions obtained with model-predicted responses.

### Identification of RGC types

To identify functionally-distinct cell types in each preparation, the retina was first visually stimulated with a white noise stimulus while recording RGC responses using a custom high-density 512-electrode array [13]. The spike-triggered average (STA) stimulus was computed for each cell to summarize its light response properties [1]. Clusters of light response properties (receptive field diameter and STA time course) were used to separate and identify specific cell types, as described previously [14–16]. In each of the four recordings examined, nearly complete populations of ON parasol, OFF parasol, ON midget and OFF midget cells were recorded. Other cell types were often recorded [15,16] but were not considered further.

### Application of CNN and LN model to ON and OFF parasol and midget cells

CNN-based models have been developed and applied to RGC recordings in several species, capturing with high accuracy responses to white noise and natural images [6,8,11,12]. To test how accurately such a model can capture the responses of the numerically dominant macaque and human RGC types to natural images, a model with a very similar architecture as one described previously [6,8], with additional learned cell-specific parameters (Fig. 1) was fitted to RGC responses to stimulation with images from the ImageNet database [17] jittered to simulate fixational eye movements [3,4] (see Methods).

**Figure 1.**
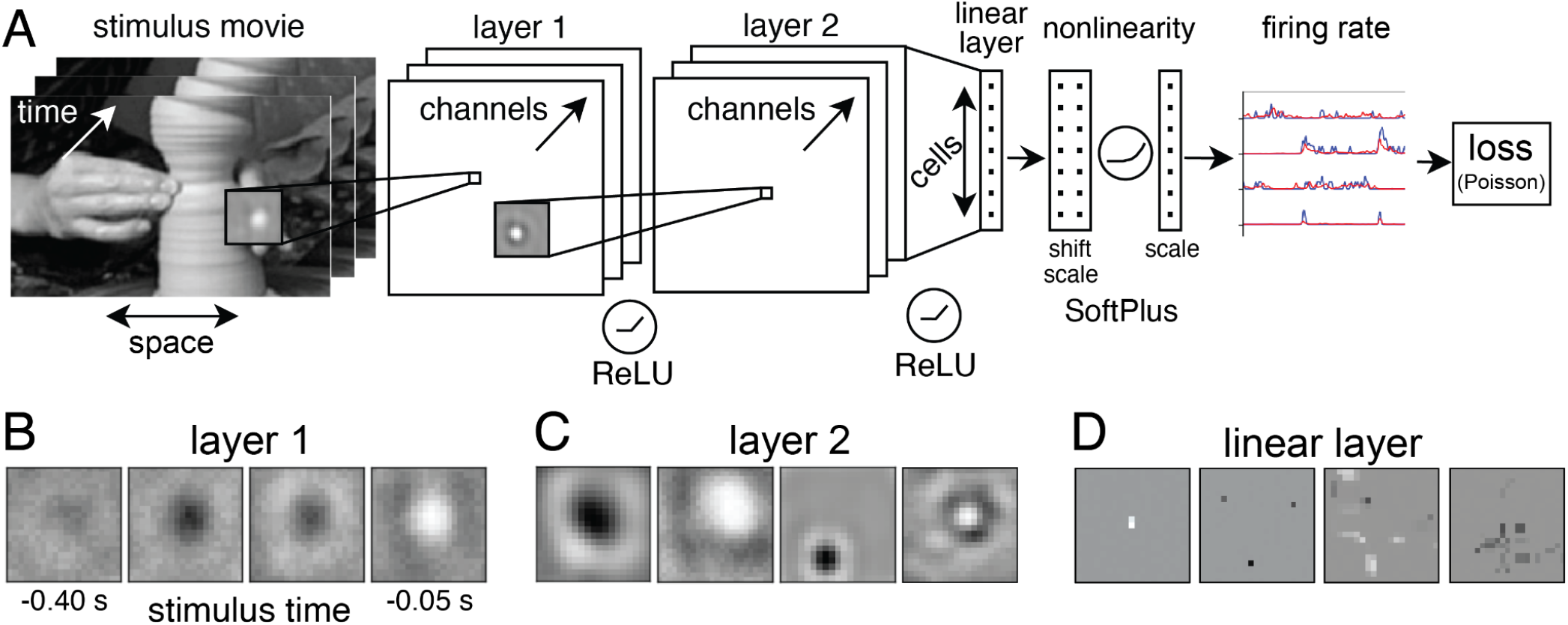
Architecture of the CNN light response model and sample learned filters. A. Illustration of the model architecture. The input at time *t* is a stimulus movie consisting of 50 frames over the time range [*t*-408 ms,*t*]. The first layer consists of eight 2D convolutions (16x16) followed by a ReLU. The second layer contains sixteen 2D convolutions (16x16) followed by a ReLU. The third layer is a fully-connected layer, which linearly maps the channels from the second convolutional layer to outputs for each cell at time *t*. The output for each cell is then shifted and scaled by learned parameters, specific to each cell. Next, these outputs are passed through a softplus nonlinearity, and then subjected to a final scaling by a third cell-specific learned parameter, producing the response for each cell at time *t*. See Methods for a full description of the model architecture. B. Example learned layer 1 filter at several times relative to *t*. C. Four layer 2 filters, each sampling from a single layer 1 channel. Layer 1 and 2 filters often exhibited a center-surround organization D. Four examples of filters in the linear layer, each sampling from a single layer 2 channel, cropped to the receptive field area. These filters often exhibited sparse input selection.

For comparison, a LN model, implemented using the same optimization framework as was used for the CNN, was also trained on the recorded data (see Methods). For each cell type, the predicted firing rates from the CNN model more closely resembled those of the true responses than did the predicted firing rates from the LN model (Fig 2). The same was true of spikes generated using a minimum variance spike generator (Fig. 3, [18]; see Methods).

**Figure 2.**
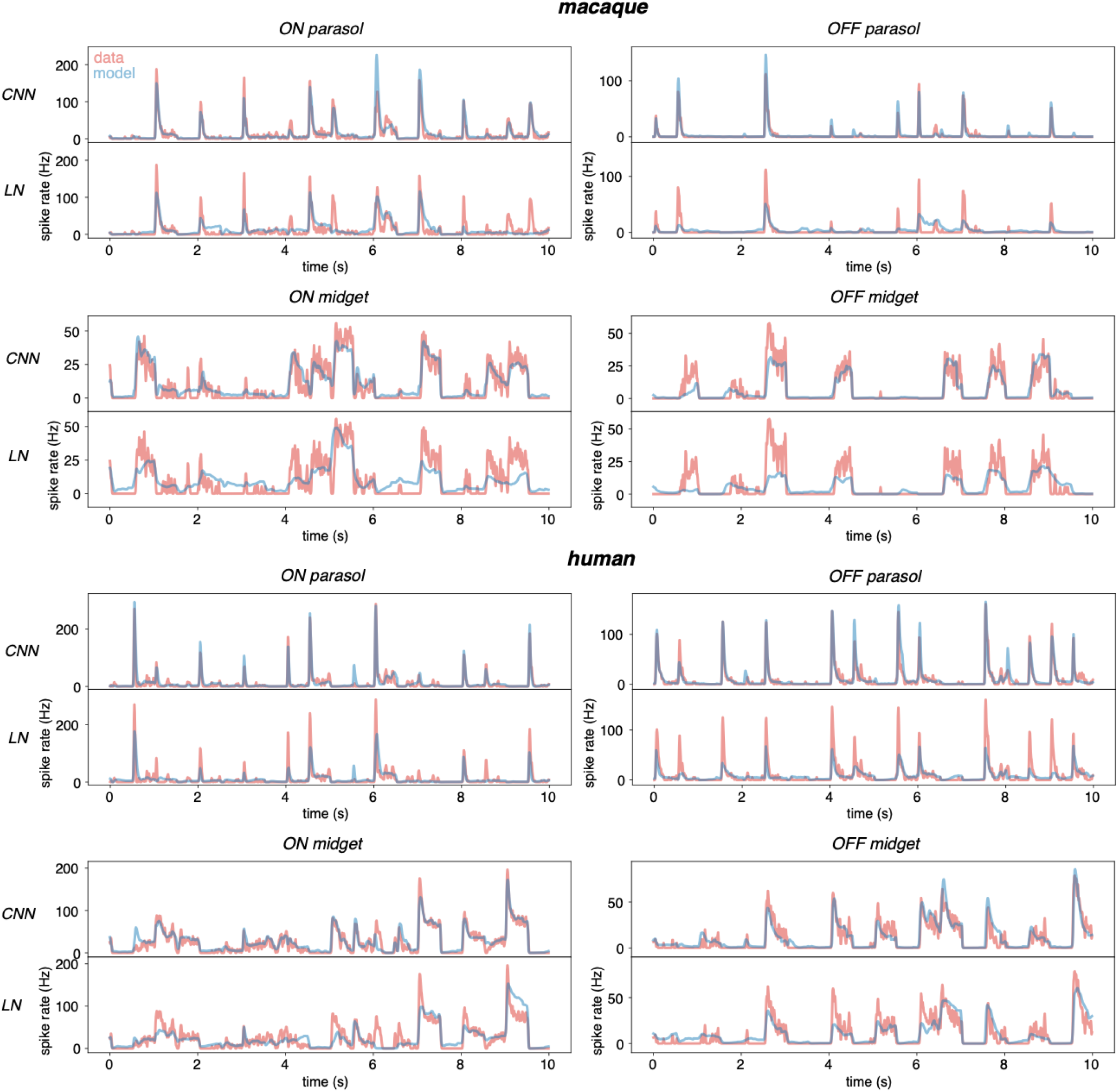
Firing rate predictions from the CNN and LN model. Measured firing rates (red) along with CNN and LN model (blue) predictions for an example ON parasol, OFF parasol, ON midget and OFF midget cells in a single preparation. The example cells selected had CNN and LN model correlation coefficients close to the median correlation across all cells of that type.

**Figure 3.**
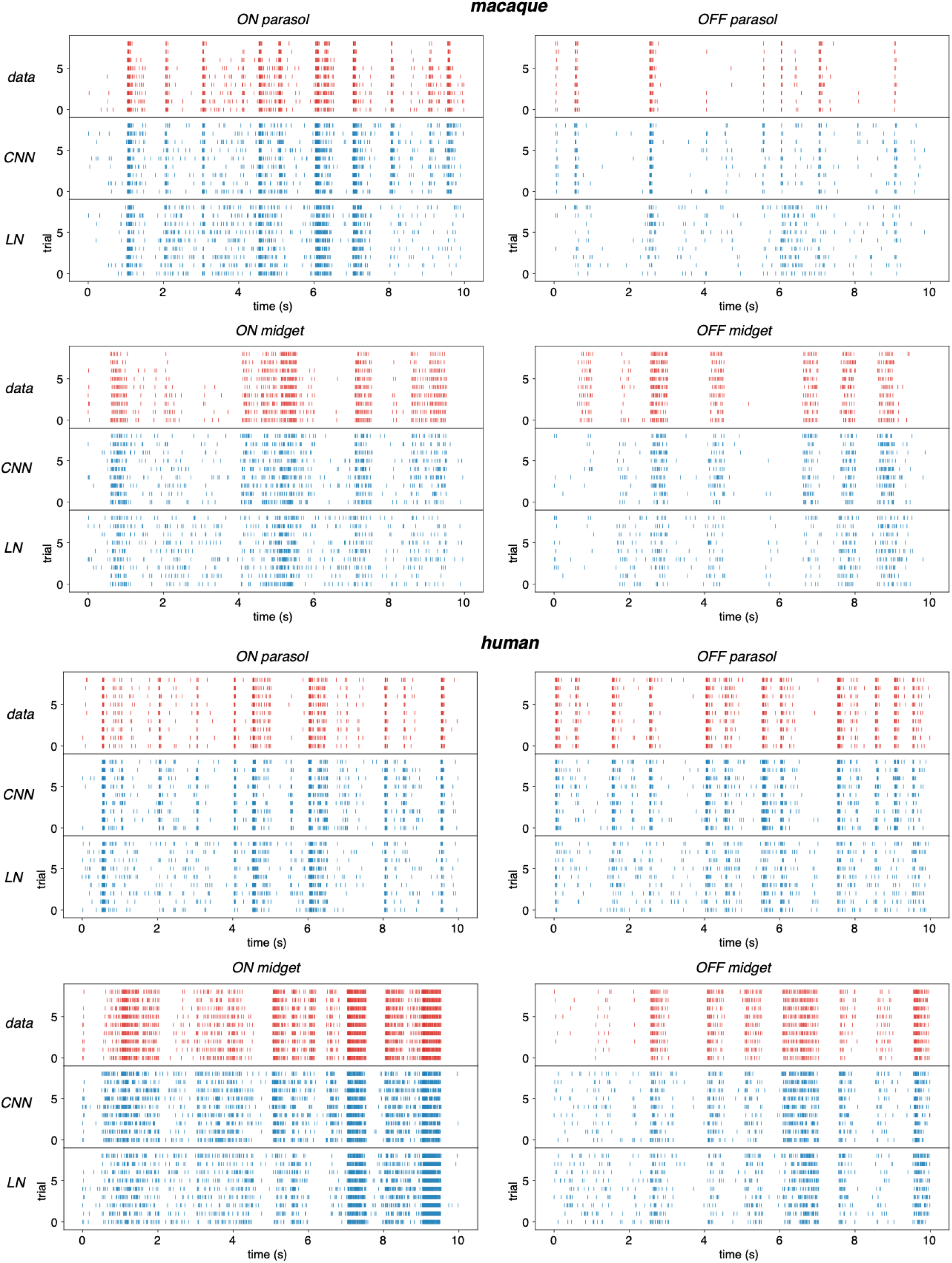
CNN and LN model predictions of spiking responses to repeated stimuli. In each sub-panel, each row represents the response of one cell to a single trial of a visual stimulus and each tick denotes a single spike. The red ticks denote the true responses and the blue ticks represent spike predictions from each model. Spike times were generated from predicted firing rates by a minimum variance spike generator (see Methods). The example cells are the same as the ones shown in Fig. 1.

The Pearson correlation coefficient was used to quantify the firing rate prediction accuracy. Across cell types and recordings, the CNN model exhibited significantly higher correlations to the recorded data than did the LN model (Fig. 4; Mann-Whitney U test, *p* < 1x10^-10^ for each cell type and preparation). The median correlation coefficient between model-predicted and measured firing rate for each cell type separately ranged from 0.74-0.91, 0.85-0.89, 0.78-0.87, 0.89-0.92 (CNN model) and 0.53-0.72, 0.66-0.73, 0.51-0.76, 0.75-0.85 (LN model), for each of the four preparations respectively (three macaque, one human). These correlation values are broadly similar, and in some cases higher, than the values reported from CNN model fits to salamander RGCs [6], and show that the model is able to capture responses to natural images of macaque RGCs with high fidelity.

**Figure 4.**
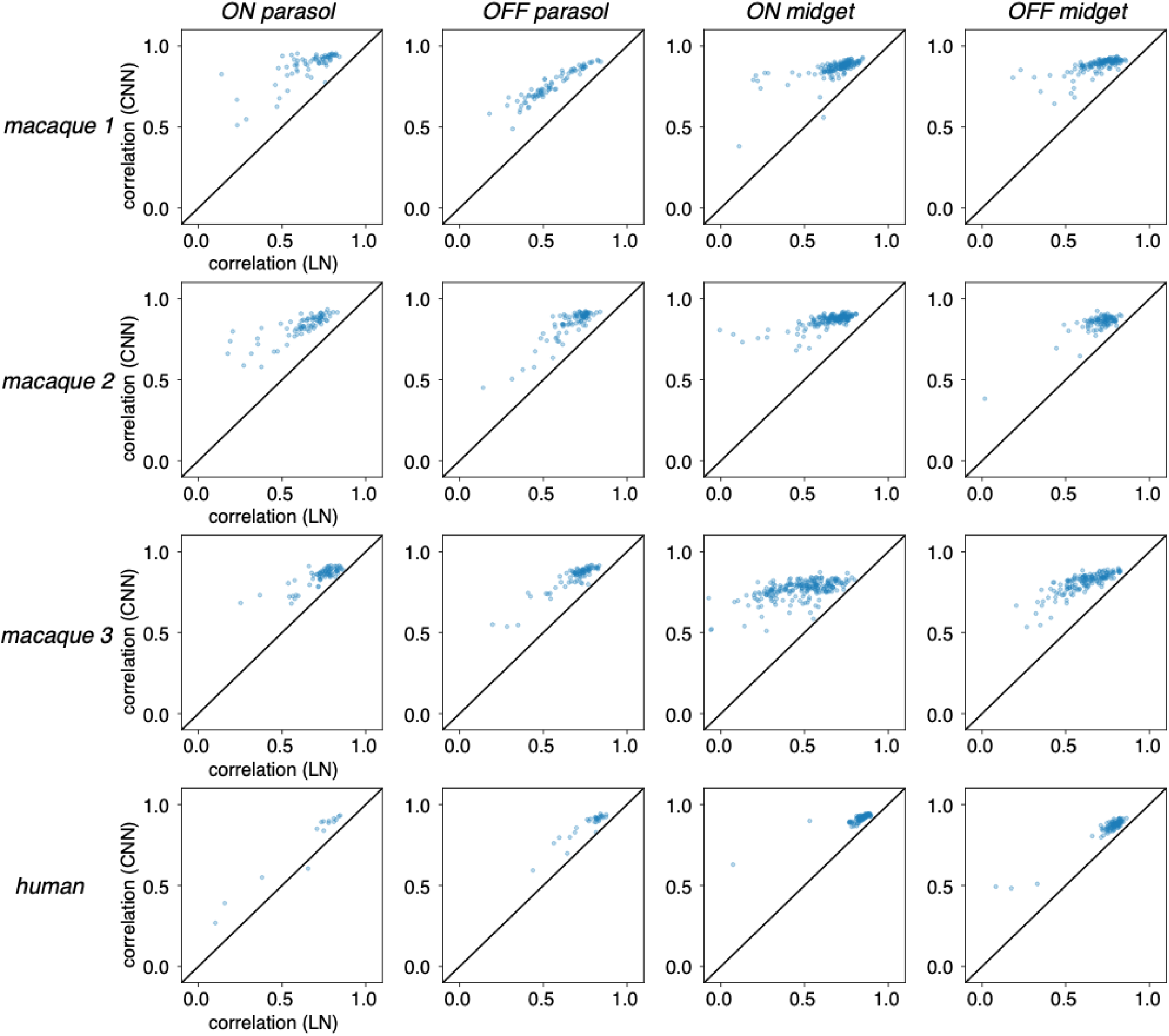
Comparison of correlation coefficients between CNN and LN model-predicted and measured firing rates. Data were accumulated across 150 test images, each displayed and jittered for 60 frames. Each point represents data from one cell, and data are shown separately for each cell type.

### Space-time linear reconstructions

Although the CNN model produced more accurate response predictions than the LN model (Figs. 2-4), whether these differences are significant for conveying visual information to the brain is unclear. To more intuitively understand the aspects of the visual neural code that are more faithfully captured by the CNN model, a spatiotemporal linear reconstruction approach was used ([19], see Methods).

Reconstructions of the stimuli from both the CNN and LN model-generated responses closely resembled those from the recorded data (Fig. 5). Image quality was quantified using the Pearson correlation coefficient and SSIM [20] between the reconstructions from the true and model-predicted responses. Images reconstructed from the CNN model-generated responses were modestly but significantly (Mann Whitney U test, *p <* 1x10^-10^) more similar to images reconstructed from the true responses than those reconstructed from the LN model (Fig. 6). Although the difference in reconstructed image quality is less striking than the difference in response prediction accuracy (Figs. 4,6, see Discussion), these findings suggest that the CNN model responses, compared to those from the LN model, capture a larger amount of the visual information that is carried by RGC spike trains.

**Figure 5.**
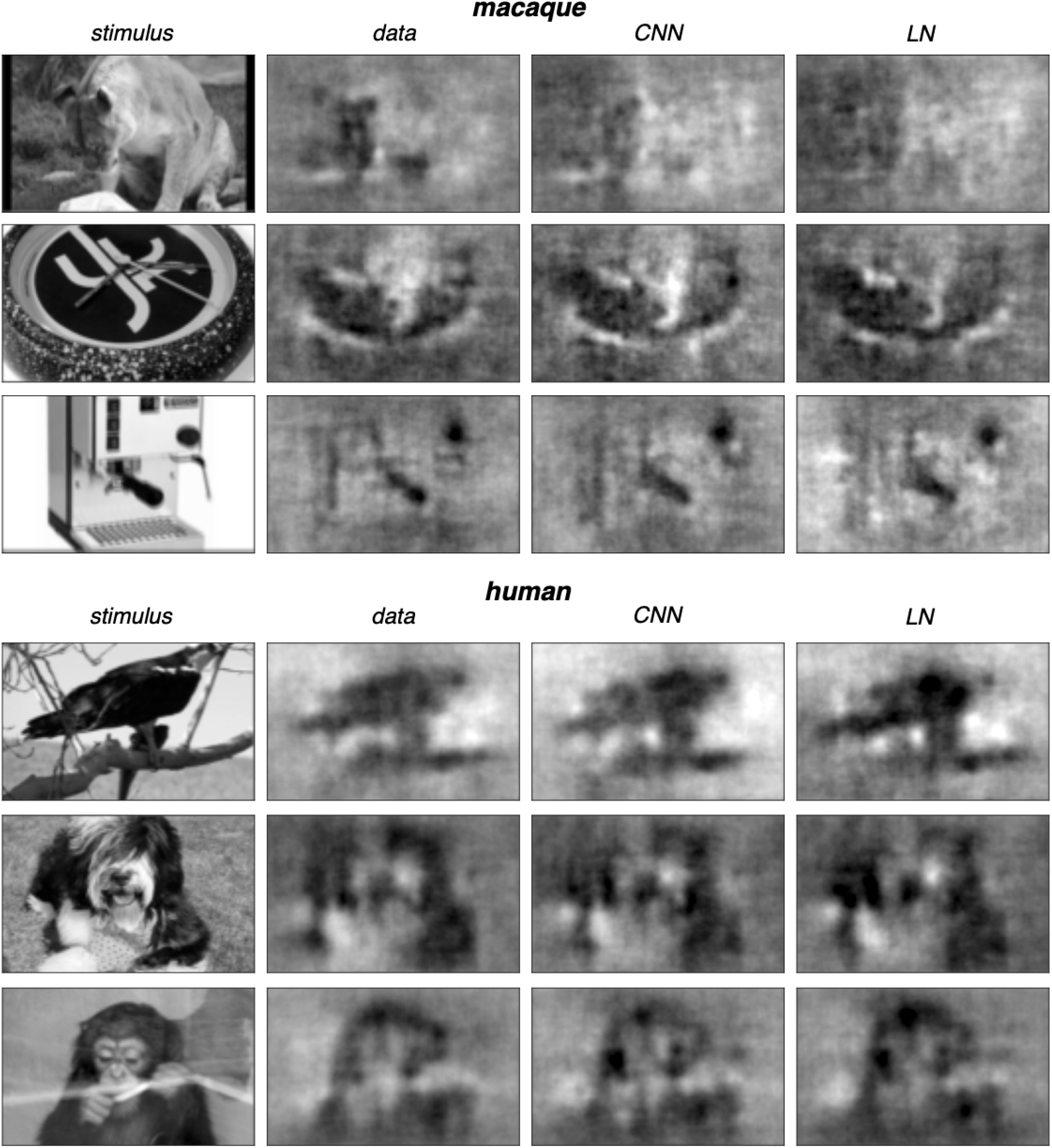
Linearly reconstructed images obtained from true and model-predicted spike trains from a single macaque and a single human preparation. Responses from ON parasol, OFF parasol, ON midget and OFF midget cells are used for each example image. The displayed image is the sum of the first 30 frames of the reconstructed image sequence (see Methods). The images shown were selected such that the difference in correlation between the 30 frames in the CNN and LN model-generated reconstructions was similar to that of the difference between the respective medians calculated across all frames.

**Figure 6.**
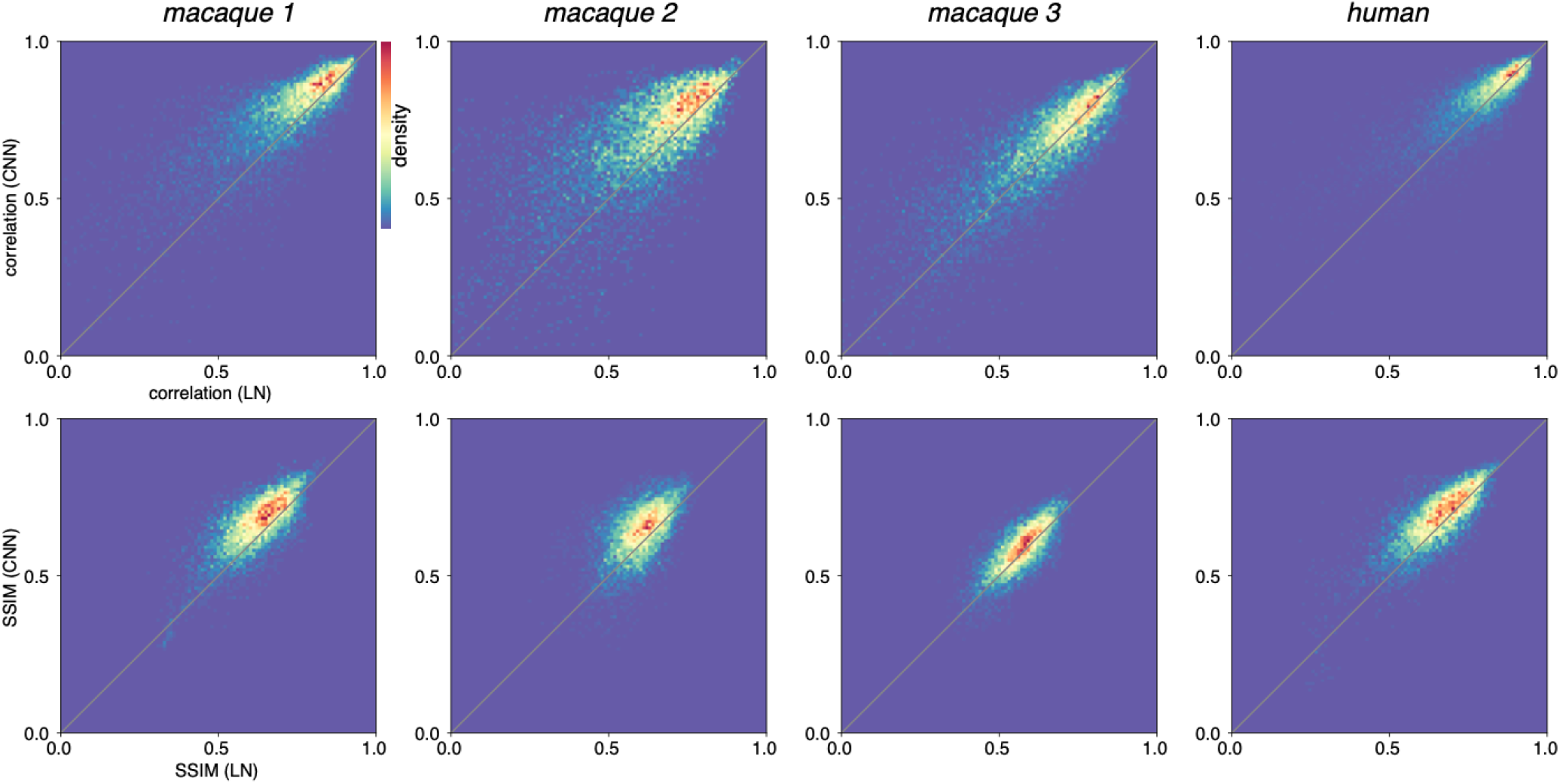
Two-dimensional histograms of reconstruction error (Pearson correlation coefficient and SSIM) accumulated across stimulus frames not used in model training, for four retinal preparations. Reconstruction was performed for 150 images jittered spatially over a duration of 60 frames each, resulting in 9000 frames linearly reconstructed (see Methods). Each reconstruction was performed using all four RGC types. For visualization, outlier values less than 0 were ignored. Across all panels, the number of excluded values per panel ranged from 34-102.

## Discussion

We evaluated the performance of a CNN-based model of the responses to natural images of the four major RGC types in macaque and human retinas. Model predictions were overall very similar to recorded responses, and the CNN-based model outperformed the LN model. An image reconstruction approach revealed that the CNN model-generated responses also more closely captured important features of the visual neural code.

Several extensions to this work could be the focus of future investigations:

- Modeling responses of additional cell types, such as small bistratified cells [16,21] and smooth monostratified cells [15,22];
- Testing whether a CNN-based model can be used to identify cell types [11] that are not easily classified by reverse correlation with white noise;
- Testing if a CNN-based model can reproduce specific ethologically relevant nonlinear computations in the retina, as was observed in the salamander retina [6]. For example, future work could focus on whether such a model can predict motion sensitivity [23–26] and direction selectivity [27,28] of RGCs in the primate retina;
- Modeling responses to continuous natural movies, potentially with varying contrast to engage gain control mechanisms [10];
- Testing to what extent model performance generalizes and how RGC receptive field properties vary across stimulus classes (i.e. natural images and white noise; [12]);
- Expanding on the linear reconstruction approach, which revealed only modest differences between the CNN and LN models despite large differences in response prediction accuracy (Figs. 4,6). The reconstruction method could be enhanced by incorporating image priors from pretrained denoisers [29] or by applying neural network approaches to either denoise the linear reconstruction [30] or decode the image directly from the responses [7,31].

As studies of visual signaling by the primate retina continue to focus on more complex and ethologically-relevant visual stimuli, the need for more sophisticated encoding models will likely increase. CNN-based models provide more accurate responses to natural scenes than the commonly used LN model, and will likely prove useful for future investigations of all stages of visual processing in primates.

## Methods

### Experimental procedures

Human eyes were obtained from a brain dead donor (33 year-old Hispanic female) through Donor Network West (San Ramon, CA). Macaque eyes were obtained from three terminally anesthetized male macaque monkeys (*Macaca mulatta, Macaca fascicularis*) used by other researchers, in accordance with Institutional Animal Care and Use Committee guidelines. After enucleation, the eye was hemisected in room lighting and the anterior portion of the eye and vitreous humor were removed. The posterior portion of the eye was stored in oxygenated, bicarbonate-buffered Ames’ solution (Sigma-Aldrich) at 31–33°C.

In dim lighting, small (2x2 mm) segments of retina with the attached retinal pigment epithelium (RPE) and choroid were separated from the sclera, and the choroid was thinned. The retina with the RPE attached was then placed retinal ganglion cell (RGC) side down on a custom high-density multielectrode array and the retina was pressed from the photoreceptor side against the array with a transparent dialysis membrane. A total of four retinal preparations (three macaque, one human) were analyzed from the peripheral retina at temporal equivalent eccentricities 32°, 40°, 46° (macaque) and 46.5° (human) [32]. Once on the array, the retina was superfused with oxygenated, bicarbonate-buffered Ames’ solution (32-35°C).

### Visual stimulation and recording

The recordings were obtained using a custom multielectrode array system with 512 electrodes (60 μm pitch, 5-15 μm inter-electrode spacing), covering an area of 1.7 mm^2^ [13]. Raw analog signals from each channel were amplified, band pass filtered (43-5000 Hz), multiplexed, digitized (20,000 Hz) and stored for off-line analysis [13]. KiloSort 2 was applied to the raw recordings to identify and separate spikes from individual RGCs [33].

Visual stimuli were delivered using a gamma-corrected CRT display refreshing at 120 Hz. Each pixel on the display was ∼5.5 μm on a side when imaged on the photoreceptor layer of the retina. White noise stimuli (pixel sizes ∼44 and ∼88 μm, refresh interval 16.67 ms) were used to identify basic light response properties of cells [1] and classify the major RGC types (see below). Natural images from the Imagenet database (dimensions 160x320, pixel size ∼11 μm, refresh interval 8.33 ms) were displayed for 500 ms, and jittered randomly on each frame according to a 2D Brownian motion (diffusion constant 10 μm^2^/frame) to emulate fixational eye movements [3,4]. A total of 5000 distinct Images were presented in sequence without gaps, interleaved with a repeating set of 150 distinct images that were each repeated 9 times.

### Cell type classification

To classify functionally-distinct cell types based on light response properties, the spike-triggered average (STA) stimulus was computed for each cell [1,32]. The receptive field (RF) of each cell was estimated by determining the pixels within the STA significantly above the noise level and fitting a 2D Gaussian to this region [15,34]. The time course of the STA was calculated by summing the three primaries of the pixels within the RF. Clusters in a 2D space formed by RF diameter and the first principal component of the time course were used to identify distinct cell types [14–16]. Cells with measurable light responses were grouped into known functional types. Typically, nearly-complete populations of ON parasol, OFF parasol, ON midget and OFF midget cells, as well as additional cell types [15], were revealed. Cells with weak to absent light responses, containing obvious contamination resulting from spike sorting errors or electrical noise events (by examining the electrical image [35] and the spike train autocorrelation) and duplicate cells were excluded from analysis and not considered further. After this classification and curation, ON parasol, OFF parasol, ON midget and OFF midget cells were considered for analysis. For each of the preparations, the following number of cells of each type were included (ON parasol, OFF parasol, ON midget, OFF midget): 66, 79,149, 134 (macaque 1); 76, 87, 169, 94 (macaque 2); 75, 79, 218, 152 (macaque 3); 15, 31, 86, 85 (human).

### CNN light response model

To capture RGC responses to natural images, a convolutional neural network (CNN) model was trained on responses to the jittered natural images [6,8]. Spike times recorded from each cell were binned with 8.33 ms precision and smoothed with a Gaussian kernel (σ = 8.33 ms) to produce an estimate of firing rate over time. The architecture of the CNN model was similar to a model described previously [6,8]. Briefly, the model has two 2D convolutional layers, a fully-connected layer and a final output nonlinearity (see below for detail). The images used for model training were downsampled 2x to a grid of size 80x160 and the mean across all images was subtracted from each image. To predict the firing rate of each cell at time *t*, the model receives as input a movie consisting of 50 stimulus frames representing the time range *t-*408 to *t* ms, and processes this input in three sequential layers.

The first layer of the model has eight 2D convolutional filters, each with a size of 50x16x16 (time x height x width). Each of these eight learned filters is convolved in space with the input movie. The resulting 50 images are then summed across time, to produce a single 2D image, defined as one channel in the output of the first layer. This operation is repeated for each of the eight learned filters, resulting in an output with eight channels (8x65x145). Batch normalization [36] is applied across all pixels in the output followed by rectification with a ReLU function (*f(x) =* max(0,*x*)).

The second layer has sixteen 2D convolutional filters, each with a size of 8x16x16 (channel x height x width). As in the first layer, each of the 16 learned filters is convolved in space with each of the eight channels emerging from the first layer. The resulting eight images in each of the 16 channels are summed, resulting in an output with 16 channels (16x50x130). Batch normalization and rectification with a ReLU are applied in the same way as in the first layer.

The third layer is a fully-connected linear layer that maps the outputs from the second layer to a vector whose dimensionality is equal to the number of cells in the recording. After the fully-connected layer, every value in the vector is shifted and scaled by learned parameters specific to each cell. Next, these values are passed through a softplus function (*f(x) =* log(1 + exp(*x*)), a smooth approximation of the ReLU function). Finally, these outputs are scaled by learned parameters specific to each cell, which produces the model firing rate of every cell at time *t*. These additional cell-specific parameters aid in more flexibly capturing potentially unique nonlinearities that could be present in different cells or cell types.

The model was trained on responses of nearly complete populations of ON parasol, OFF parasol, ON midget and OFF midget cells to a non-repeating set of 4850 images. Responses to a distinct, non-repeating set of 150 images were used for model validation and responses to a third distinct set of 150 images repeated nine times were used for final evaluation. Mini-batch gradient descent (batch size 32, learning rate 0.0001, 10 training epochs guided by the validation set) was performed using the Adam optimizer [37] on pairs of natural images and simultaneously recorded RGC responses. Model parameters were randomly initialized and optimized to minimize Poisson negative log-likelihood loss. To prevent overfitting, in both of the convolutional layers, L2 regularization was applied to the filters and Gaussian noise was added to the layer outputs [8] after batch normalization and before application of the ReLU. L1 regularization was applied to the linear weights enforcing sparse feature selection from the output of the second layer (Fig. 1D). The L1 and L2 penalty weights were selected based on performance on the validation set.

All model training software was developed using PyTorch and is publically available on GitHub: https://github.com/Chichilnisky-Lab/primate-cnn-model.

### Linear-nonlinear (LN) cascade model

To model RGC responses to natural images with a simpler architecture, a linear-nonlinear (LN) cascade model was trained on responses to natural images using the same optimization method used for training the CNN model. Five training epochs were used for two preparations and 10 training epochs for the other two preparations, guided by the validation set. The visual stimulus input used by the LN model is a stimulus movie (50 frames from time *t* to *t*-408 ms). The LN model consists of a fully-connected linear layer followed by a softplus nonlinearity with fitted scaling and offset parameters as in the CNN model. To prevent overfitting, L1 regularization was applied to the weights of the fully-connected linear layer.

### Model fit variability

For both models, random initialization of parameters and stochastic optimization can generate variable model performance across training instances. In the Results, we analyzed only a single CNN or LN model training instance for each preparation. To understand the potential impact of training variability across model instances, we computed the firing rate prediction correlation coefficients for all cells in a single macaque retina (*macaque 1*) across five CNN and LN model training instances. We then took the signed difference between correlation coefficients obtained from pairs of CNN and LN model instances (*corr*_*CNN*_ *-corr*_*LN*_), yielding five signed difference vectors, each containing an entry for every cell. Finally, we computed pairwise correlations between these signed difference vectors. The mean of these values was 0.95, revealing that despite non-deterministic model performance, different training instances result in very similar CNN versus LN model performance.

### Firing rate analysis

For each model, firing rate predictions of each cell were generated on a set of 150 repeating images (9 repeats total) unseen by the model. To estimate the true firing rate for each cell, smoothed spike counts were averaged over responses to each repeated image. To quantify model performance, the pointwise Pearson correlation coefficient between the model generated and true firing rate was calculated for each cell.

### Spike train analysis

For each model, the ability to capture spike train structure was explored by adding a spike generator to the model. Classically, spike trains are modeled as a Poisson process, where the rate parameter represents the instantaneous spike probability. However, in practice, RGC spiking exhibits sub-Poisson count variability, i.e. the variance in spike counts is much smaller than the mean. Therefore, a *minimum variance* model of spiking was used, which more accurately captures RGC spiking statistics than a Poisson process does. In the minimum variance spike generator, for the time bin centered at time *t*, if the the firing probability associated with that bin is given by *f*(*t*), then ⌊*f*(*t*) ⌋ spikes are generated with probability *p* =⌊*f*(*t*) ⌋ −*f*(*t*), and ⌊*f*(*t*) ⌋ spikes are generated with probability 1−*p*[18]. This Bernoulli process minimizes the spike count variance for a given spike count mean and more accurately captures RGC spiking statistics than does a Poisson process [18].

### Image reconstruction analysis

To more intuitively understand how well each model captures the visual neural code of each RGC, a linear spatiotemporal image reconstruction approach was applied, as described previously [4,19]. For quantitative comparison, SSIM [20] and the Pearson correlation coefficient were computed between the reconstruction obtained from data and that decoded from model generated responses, using every frame from a single trial of images in the test set. For display purposes, the sum of the first 30 jittered frames of each image are shown, based on the typical duration of each cell’s spatiotemporal reconstruction filter. Although the frames of each image in the stimulus were jittered, taking the sum without jitter correction results in minimal blurring because of the spatial downsampling that was performed and because the degree of jitter is not large relative to spatial structure of the images.

## Acknowledgements

The human eye was provided by Donor Network West (San Ramon, CA). We are thankful for the cooperation of Donor Network West and all of the organ and tissue donors and their families, for giving the gift of life and the gift of knowledge, by their generous donations. We thank T. Moore, SRI International and The California National Primate Research Center for access to primate retinas; S. Baccus, J. Melander, A. Nayebi, S. Idrees, J. Zylberberg, F. Rieke and G. Field for useful discussions and comments on the manuscript; R. Samarakoon and S. Kachiguine for technical assistance. This research was supported by NIMH T32MH020016, NEI F31EY033636, Fondation Bertarelli, the Stanford Neurosciences Graduate Program (ARG), a donation from John Chen (AML), Stanford Medicine Discovery Innovation Award, Stanford Bio-X Interdisciplinary Initiatives Program Seed Grant, NEI R01 EY033870, NEI R01 EY029247, NEI R01 EY032900, and NEI P30-EY019005 (EJC).

